# Environmental differences impact *Acinetobacter baumannii* phage isolation and infectivity

**DOI:** 10.1101/2024.07.10.602838

**Authors:** Ellinor O Alseth, Carli Roush, Iris Irby, Mykhailo Kopylov, Daija Bobe, Kristy Nguyen, Huaijin Xu, Anton V Bryksin, Philip N Rather

## Abstract

With the global rise of antimicrobial resistance, phage therapy is increasingly re-gaining traction as a strategy to treat bacterial infections. For phage therapy to be successful however, we first need to isolate appropriate candidate phages for both clinical and experimental research*. Acinetobacter baumannii* is an opportunistic pathogen known for its ability to rapidly evolve resistance to antibiotics, making it a prime target for phage therapy. Yet phage isolation is often hampered by *A. baumannii*’s ability to rapidly switch between capsular states. Here, we report the discovery and structural characterisation of a novel lytic phage, Mystique. This phage was initially isolated against the wild-type AB5075: a commonly used clinical model strain against which no phage has previously been readily available for the capsulated form. When screening Mystique on 103 highly diverse isolates of *A. baumannii*, we found that it has a broad host range, being able to infect 85.4% of all tested strains when tested on bacterial lawns – a host range which expanded to 91.3% when tested in liquid culture. This variation between solid and liquid environments on phage infectivity was also observed for several other phages in our collection that were assumed unable to infect AB5075, and capsule negative mutants that initially seemed completely resistant to Mystique proved susceptible when assayed in liquid. Overall, through the discovery of a novel phage we demonstrate how environmental differences can drastically impact phage infectivity with important consequences for phage isolation and characterisation efforts.

**Author summary:** Bacterial infections caused by *Acinetobacter baumannii* are a major global health concern due to high antibiotic resistance, earning it a critical priority pathogen ranking by the WHO. Phage therapy is resurging as a treatment option, with some success against *A. baumannii*. However, the wild-type clinical model strain used to assess new therapies lacks an available phage, and isolating phages for *A. baumannii* is challenging due to its complex capsule. Here, we report the discovery of a novel lytic phage, Mystique, which exhibits a broad host range, infecting 94 out of 103 tested *A. baumannii* strains. We conducted genomic sequencing and structural analysis to fully characterise Mystique. Additionally, we found that the testing environment significantly impacts results; some phages that do not form plaques on bacterial lawns can still infect and amplify in liquid cultures of the same strain. Moreover, mutants resistant to Mystique based on plaque assays were susceptible in liquid culture assays. This work underscores the necessity of a multifaceted approach for phage isolation and characterisation, as traditional phage assays may not be sufficient for studying bacteria-phage dynamics in certain bacteria such as *A. baumannii*.

## Introduction

*Acinetobacter baumannii* is an increasingly antibiotic resistant and virulent bacterium, known to cause severe nosocomial infections [1–4]. With an estimated 63% of isolates in the United States being considered multidrug resistant [5], infections due to *A. baumannii* are difficult to overcome and often fatal [6]. An additional challenge is *A. baumannii’*s ability to contaminate and persist in healthcare facilities, such as in laminar flow systems [7], on care and medical equipment [8–10], and on other surfaces like curtains, doors handles, and keyboards [10,11]. It is largely due to this challenge of preventing and treating *A. baumannii* infections that the therapeutic application of bacteriophages (phages, *i*.*e*. viruses that infect bacteria) is increasingly being considered and – usually as a last resort – applied [12,13].

Microbial model systems play a crucial role in advancing science across various biological disciplines [14,15], and are therefore important to increase our understanding of bacteria-phage dynamics in ways that can help improve the efficacy of phage therapy. For instance, model systems provide researchers with a highly controlled environment for testing experimental predictions that may elucidate fundamental principles underlying bacteria-phage interactions, such as mechanisms of phage infection [16], phage resistance [17], and coevolutionary dynamics [18]. Additionally, microbial model systems serve as helpful tools when exploring the application of phages as therapeutics and their potential consequences [19]. After all, the arms race between phage (‘predator’) and bacteria (‘prey’) is fast paced, with a strong selection pressure for the bacteria being targeted to evolve phage resistance both *in vitro* and *in vivo* [20]. Well-characterised model systems for studying bacteria-phage interactions have been developed for other opportunistic pathogens such as *Escherichia coli* [21,22] and *Pseudomonas aeruginosa* [23,24], yet for some clinically relevant strains of *A. baumannii* there are no phages available. This is the case for the wild-type clinical isolate and model strain AB5075 in particular. AB5075 is a highly virulent strain of *A. baumannii* that is commonly used in various animal models to study pathogenesis, host-pathogen interactions, and to assess potential new treatments [25].

In an otherwise genetically homogenous population, AB5075 and other *A. baumannii* strains exhibit phenotypic heterogeneity by rapidly switching between virulent opaque (VIR-O) and avirulent translucent (AV-T) colonies [26–28]. This phenotypic switch is associated with changes in capsule thickness, with AV-T cells exhibiting a twofold decrease in capsule thickness compared to VIR-O cells [27]. While the switching frequency is at ∼4-13% over 24 hours for a single propagated colony, this rate is potentially affected by the selection pressure imposed by the presence of phages targeting one state but not the other. In other words, the process of isolating novel phages can incur an increased selection pressure for rapid capsule modulation, at a rate incompatible with commonly applied methods of phage isolation. For *A. baumannii*, many phages have been found to select for reduced capsule production [29,30], but most phage isolation attempts for AB5075 specifically have resulted in phages targeting capsule negative mutants [31,32], but not the VIR-O state or the wild-type. It is therefore possible that a thicker capsule might directly block phage absorption; a characteristic found in other bacteria with similar capsule properties [33],

Here, we report our successful approach to phage isolation for AB5075, resulting in the isolation of Mystique: a novel lytic phage against *A. baumannii*. In addition to infecting AB5075, Mystique can readily infect 91.3% of a diverse set (*n* = 103, including AB5075) of clinical *A. baumannii* isolates. This makes for a remarkably broad host range, especially as the strains Mystique was tested on are specifically meant to represent the full genetic diversity of *A. baumannii* as a species [34]. To isolate phage Mystique, we combined raw sewage with a cocktail of known phages that can infect but not plaque on AB5075, to limit resistance evolution occurring during the isolation process. The isolation process for this phage also revealed how a structured environment can result in false negative results when assessing phage infectivity, as we discovered multiple phages in our collection were able to infect AB5057 in liquid culture but not on a bacterial lawn. Additionally, capsule mutants of AB5075 that would traditionally be classified as resistant to Mystique based on plaque assays proved susceptible to the phage in liquid cultures. This has important implications for any phage isolation attempt for *A. baumannii* and other bacteria with similar capsule properties, seeing as most phage assays are performed on bacterial lawns: a method that has not seen much change since the discovery of phages in the early 1900s [35,36].

## Results

### Isolation of novel *A. baumannii* phage Mystique

There is currently no lytic phage readily available against the wild-type clinical model strain of *A. baumannii*: AB5075, and yet a well characterised phage is essential to study clinically relevant bacteria-phage dynamics. To solve this issue, we isolated a novel lytic phage, named Mystique, from local sewage water in Atlanta, USA, using AB5075 as the bacterial target for phage infection (Fig 1). To aid in the isolation attempt, other phages were added to the mixture, to limit the likelihood of resistance evolution (Fig 1A and Fig 2).

**Figure 1.**
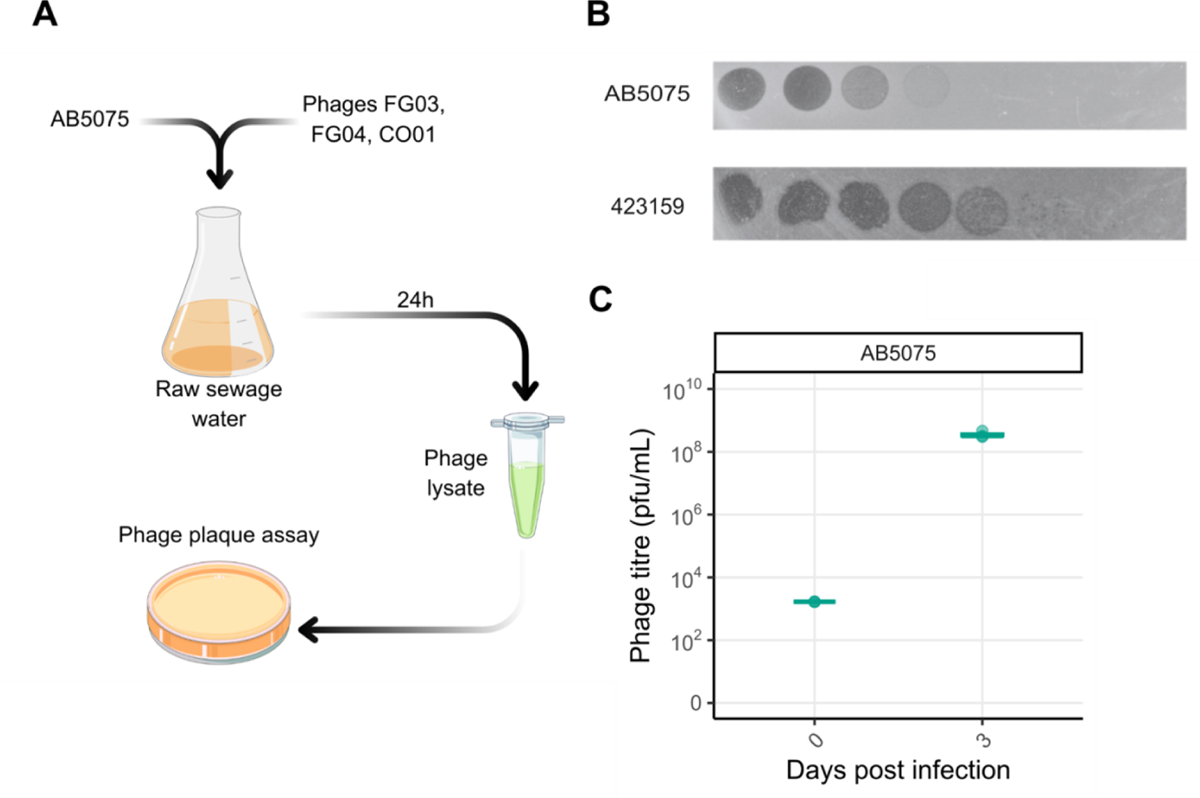
Mystique is a novel lytic phage targeting *A. baumannii* AB5075. **A** Illustration of the phage isolation process. **B** Serial dilution of Mystique phage lysate after isolation and purification (always using AB5075 as host for phage amplification) to ensure the presence of only one phage, pipetted onto lawns of *A. baumannii* strains AB5075 and 423159. **C** Mystique phage titres after three days of co-inoculation and daily passaging with AB5075.

**Figure 2.**
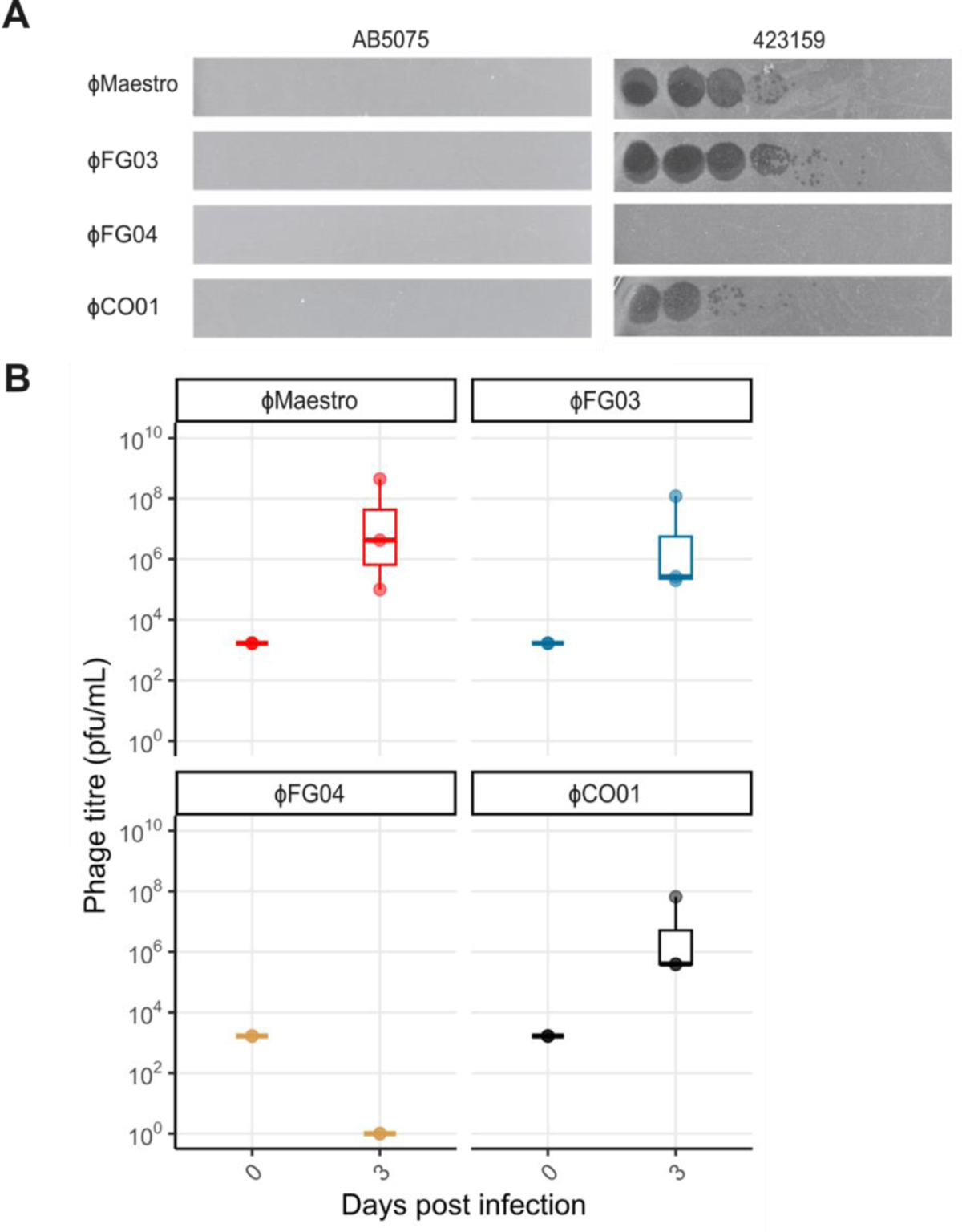
Some phages can infect in liquid culture but not on a bacterial lawn of **the same host strain. A** Bacterial lawns of AB5075 and 423159 with serial dilutions of phages Maestro, FG03, FG04, and CO01, illustrating how none of them plaque on AB5075 but three do on 423159, which was used as an indicator for which strains might be able to infect AB5075 in liquid cultures. **B** In liquid culture on the other hand, Maestro, FG03, and CO01 are all able to infect AB5075 (phage plaques were counted on lawns of *A. baumannii* strain 423159 as they do not plaque on AB5075 as shown in A). FG04 cannot infect AB5075 on a lawn or in liquid culture.

While Mystique does cause bacterial clearance on a lawn of AB5075, it does not form individual phage plaques, which made it difficult to verify the presence of a single phage (Fig 1B). However, when pipetting a serial dilution of the sewage lysate on various strains of *A. baumannii*, phage plaques were observed on *A. baumannii* MRSN strain 423159 [34] (Fig 1B), from which an individual plaque was picked and purified three times in liquid cultures of AB5075 to ensure the isolation of one individual phage. Plaque assays on 423159 were also used to assess Mystique’s infectivity in liquid cultures of AB5075, which revealed the phage’s ability to infect, amplify, and remain in the population over the course of three days (Fig 1C).

### The phage isolation process revealed how some phages can only infect in liquid culture

Initially, the isolation of phage Mystique seemed to have been in vain, as no phage plaques were observed on a lawn of AB5075 after our first isolation attempt. We hypothesised that the rapid phenotypic switching between VIR-O and AV-T states, as a potential strategy to become phage resistant, could be a significant hindrance in the initial phage amplification step. In an attempt to limit the evolution of phage resistance, we then reasoned that the addition of other known phages against *A. baumannii* could limit rapid capsule switching and the emergence of phage resistant mutants.

In our phage biobank we have three phages (Maestro [20], FG03, and CO01 [30]) that also plaque on *A. baumannii* strain 423159 (Fig 2A). In particular, Maestro has been used clinically against *A. baumannii* strain TP1 as part of a cocktail including several phages that were initially isolated on AB5075 capsule negative mutants[13,20,31], indicating that this phage might also be able to infect AB5075 – but only one capsule state. To test the ability of these other phages to infect AB5075, we inoculated them with AB5075 in liquid culture for three days with daily transfers into fresh medium. FG04 [30] was included as a negative control as it does not plaque on MRSN strain 423159, and we consequently expected that it would not be able to infect AB5075 either (Fig 2A). We found that the phages Maestro, FG03, and CO01 all infect and amplify on AB5075 in liquid culture, while FG04 did not (Fig 2B). All plaque assays were performed on lawns of 423159 at three days post infection, to make plaque counting possible.

Based on these findings, we subsequently added phages Maestro, FG03, FG04, and CO01 to the sewage before inoculation with AB5075, which resulted in the isolation of phage Mystique (Fig 1). The addition of these phages to the sewage filtrate might therefore have aided in the isolation of phage Mystique, with Maestro, FG03, and CO01 constraining resistance evolution while not confusing the results due to their inability to plaque on AB5075 – unlike Mystique. FG04 was also included to further reduce the likelihood of drastic mutational changes to a potential phage surface receptor, as any such change could potentially come at the cost of making the host susceptible to this other phage.

### Mystique sequencing, annotation, and assembly

After the isolation and purification of Mystique, DNA was extracted from the phage lysate followed by Illumina and Oxford Nanopore sequencing. Once the hybrid (long and short sequencing reads) assembly of the genome was complete, Mystique was found to be a dsDNA phage closely related to two other *Acinetobacter* phages: vB_AbaS_TCUP2199 (GeneBank accession number ON323491.1 [37]) shows 96.63% identity across 97% of the Mystique genome, and EAb13 (GeneBank accession number OQ717042.1 [38]) has 84.47% identity across 8% of the Mystique genome. The genome has a GC-content of 40%, with 154 predicted genes of which 115 are annotated as hypothetical proteins, while 38 have assigned putative functions. Additionally, one tRNA gene was identified (Fig 3).

**Figure 3.**
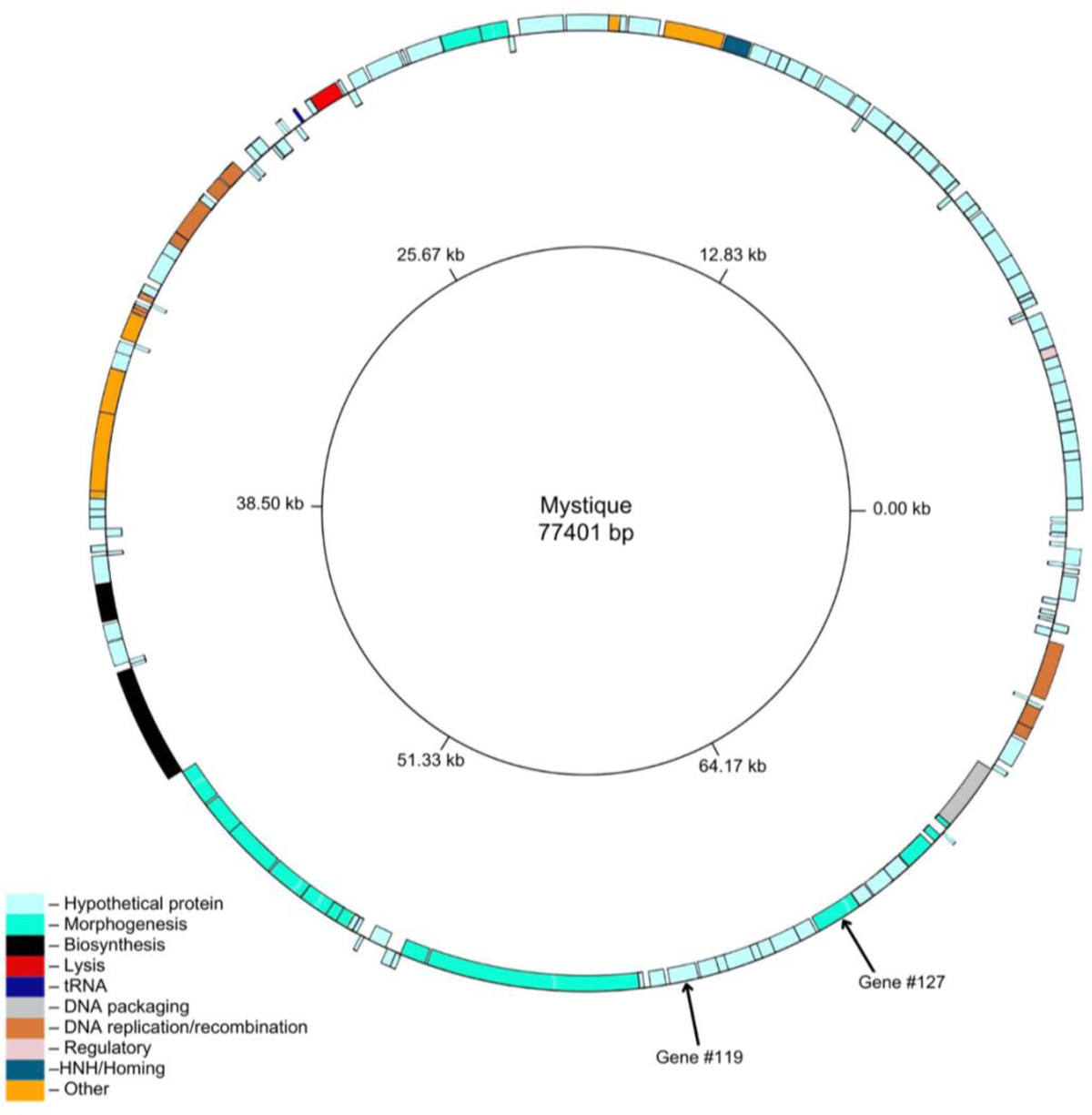
The annotated genome of Mystique. Mystique has a genome size of 77,401bp, with 154 predicted genes out of which the majority are hypothetical and 38 have putative functions. Figure generated using GenomeVx [39].

### CryoEM analysis reveals an icosahedral T=9 capsid and a long flexible tail

To gain insights into the phage morphology and structure of Mystique, we analysed the phage lysate with negative-stain Transmission Electron Microscopy (TEM) followed by cryo electron microscopy (cryoEM). Negative staining was done to verify sample viability and concentration, after which the phage was further concentrated using polyethylene glycol (PEG) precipitation [40]. CryoEM data acquisition revealed that phage particles were present in vitreous ice on CryoEM grids, yet most of the tails were detached from the heads, possibly due to the harsh conditions of PEG precipitation [41]. In total, 7200 head particles and 363,000 tail segments were picked from micrographs for further analysis. Interestingly, the initial 2D classification revealed that the Mystique head particles were in two distinct states – one empty and one full (Fig 4A and B). A total of 6000 head and 191,150 tail segments were used for final refinement, producing maps at 4.5 Å and 3.2 Å resolution respectively (Fig 4C). Additionally, independent 3D reconstruction of particles from both empty and full states (Fig 4B) produced lower resolution, but identical maps, suggesting that phage capsid structure does not depend on the presence or absence of nucleic acids (Fig 4D).

**Figure 4.**
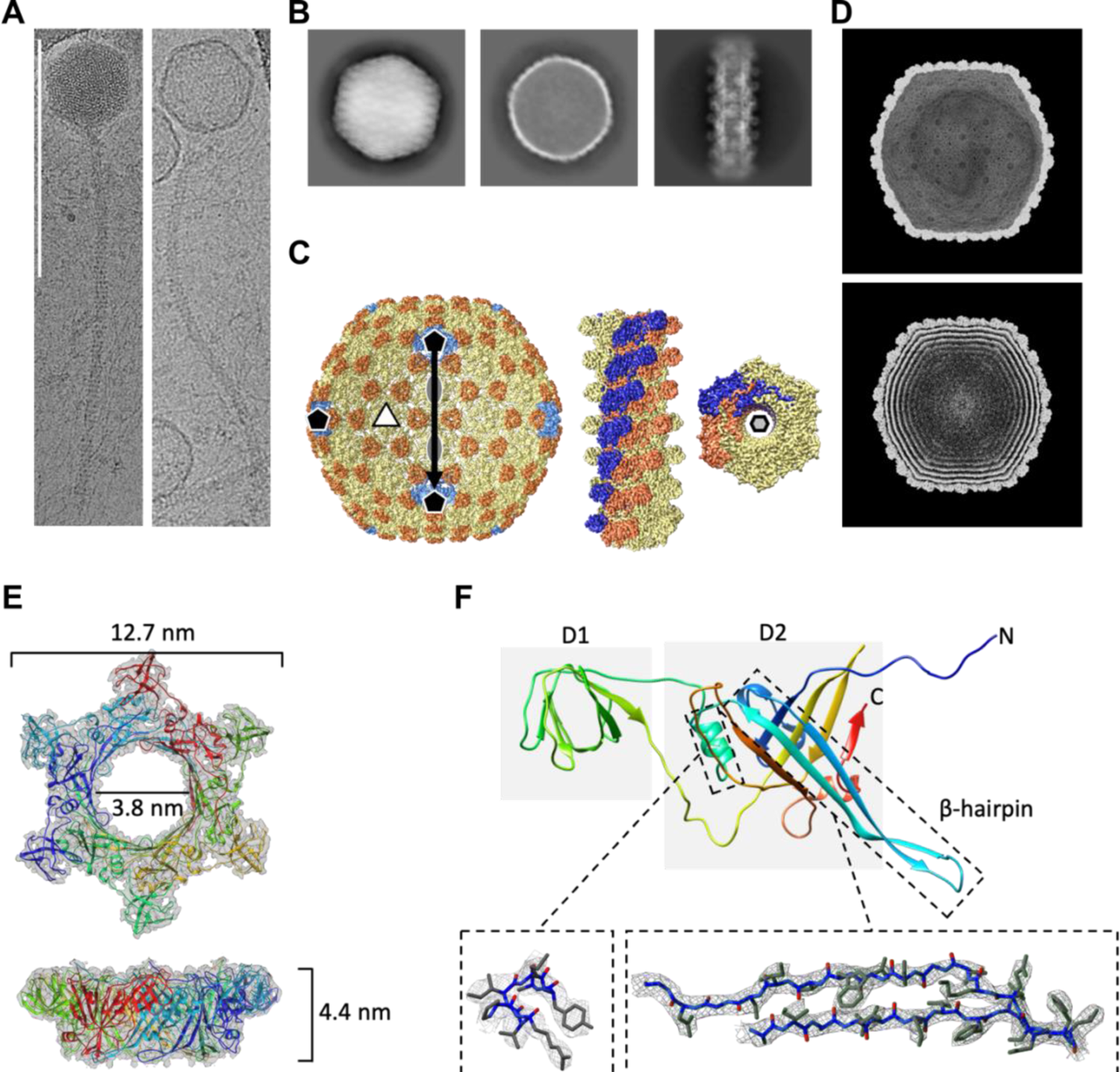
CryoEM reconstruction of bacteriophage Mystique and structural tail details showing hexameric assembly. **A** Individual phage particles from CryoEM micrographs showing bacteriophages with a full head (left) and an empty head (right). Scale bar 200 nm. **B** Selected 2D classes from data processing of phage particles (left = full head, middle = empty head, right = tail). **C** Left: 3D reconstruction (with icosahedral symmetry applied) of the Mystique head showing icosahedral symmetry axes. For this map, particles from empty (2,793) and full (3,207) heads were combined resulting in a 4.5 Å resolution reconstruction (blue = pentamers, yellow = hexamers, orange = decorating protein). Right: helical reconstruction of Mystique tail with C6 symmetry. Two adjacent individual helical strands are coloured in blue and orange. **D** Independent CryoEM reconstructions of empty (top) and full (bottom) capsids displayed at high thresholds. **E** Helical asymmetric unit of Mystique’s tail unit with a 6-fold symmetry. The final refinement had a helical twist of 17.9 degrees and a helical rise of 42 Å. **F** Tail protein monomer with two main domains and a β-hairpin. Boxes show the alpha helix and a β-hairpin fitted into a 3.2 Å map.

Through cryoEM, we further found that Mystique’s head has a icosahedral T=9 symmetry (h=3, k=0 [42]), and AlphaFold2 structure prediction of the protein encoded by Mystique gene number 127 (Fig 3) revealed an HK97 fold [43], suggesting a similar organisation to that of other phages with HK97 capsids (Fig S1A). However, the resolution of the phage heads was not sufficient to unambiguously trace a backbone model in cryoEM map density. Instead, we used rigid-body fitting to manually fit the predicted model into the map, assembling an asymmetric unit (Fig S1B) before applying icosahedral symmetry to produce a closed cage that matched the EM density (Fig S1C). Unfilled densities may belong to a yet unidentified “cement” or “decoration” protein common for bacteriophages with HK97-like folds [43].

Next, the cryoEM map of the phage tail was used for protein sequence prediction though *de novo* modelling using ModelAngelo [44]. The resulting model was then used to search through Mystique’s genome, identifying gene number 119 as the tail protein (Fig 3). The protein sequence derived from gene number 119 was further used to build a model which was refined with C6 symmetry (Fig 4E) before being used as input for Foldseek [45]. This revealed that YSD1, a phage infecting *Salmonella* [46], has a highly homologous tail structure, despite low sequence similarity (Fig S2A and B). The Mystique tail monomer is organised into two major domains: the external D1 domain and a core D2 domain. A β-hairpin of each subunit in a hexamer interacts with a preceding and two subsequent subunits, thus forming a highly interlocked assembly (Fig 4F). A third domain (D3) is absent in the Mystique tail, but present in the YSD1 tail [46], although poorly resolved. Similar to YSD1, Mystique’s tail also has a highly negatively charged lumen necessary for translocation of nucleic acid from the head to the host (Fig S2C).

### Mystique has a broad *A. baumannii* host range

With Mystique sequenced and structurally characterised, we next set out to determine its more exhaustive *A. baumannii* host range. We tested Mystique’s killing ability using plaque assays on 103 highly diverse clinical isolates of *A. baumannii*, 100 of which were from the MRSN diversity panel [34] as well as FZ21 [17] and TP1 [13], with AB5075 included as a positive control. The MRSN diversity panel in particular is meant to represent the full genetic diversity of *A. baumannii* as a species [34].

Based on these plaque assays, we found that Mystique can infect at least 88 out of the 103 strains tested (85.4%) (Supporting information 1), making it a broad host-range lytic phage. However, because we previously showed that the environment for assessing phage infectivity is significant (Fig 2), we hypothesised that Mystique might have an even broader host range than indicated by plaque assays if tested in a liquid culture rather than on a bacterial lawn. To determine this, we next inoculated Mystique in broth culture with the 15 strains it does not plaque on, with daily transfers for three days before phage titres were determined by plaque assays on 423159. In liquid culture, in addition to the 88 already confirmed, Mystique could also infect MRSN strains 334, 1171, 7153, 11816, 22112, and 337038 (Fig S3) for a total of 94 out of the 103 strains tested (91.3%). These results further highlight the importance of testing phages in liquid culture, as there was once more a large discrepancy in the observed results between environments (lawn vs liquid) used to test for phage infectivity (Fig S3).

Next, we made a phylogenetic tree and tested for a potential phylogenetic signal across strains by first assessing strains susceptible based on plaque assays (Fig 5). This gave us a D value of 0.3 (*p* = 0), which is a measure of phylogenetic signal for a binary trait, where a number closer to 1 indicates a trait evolved from random motion [47]. With susceptibility in liquid as the binary trait on the other hand, the D value decreased to 0.084 (*p* = 0.02), strongly indicating that resistance to Mystique is a non-random and heritable trait (Fig S4). There were no clear similarities between the susceptible strains’ capsule serotype (KL) or lipooligosaccharide outer core locus (OCL) that could answer why a specific strain is resistant or susceptible to Mystique (Fig 5).

**Figure 5.**
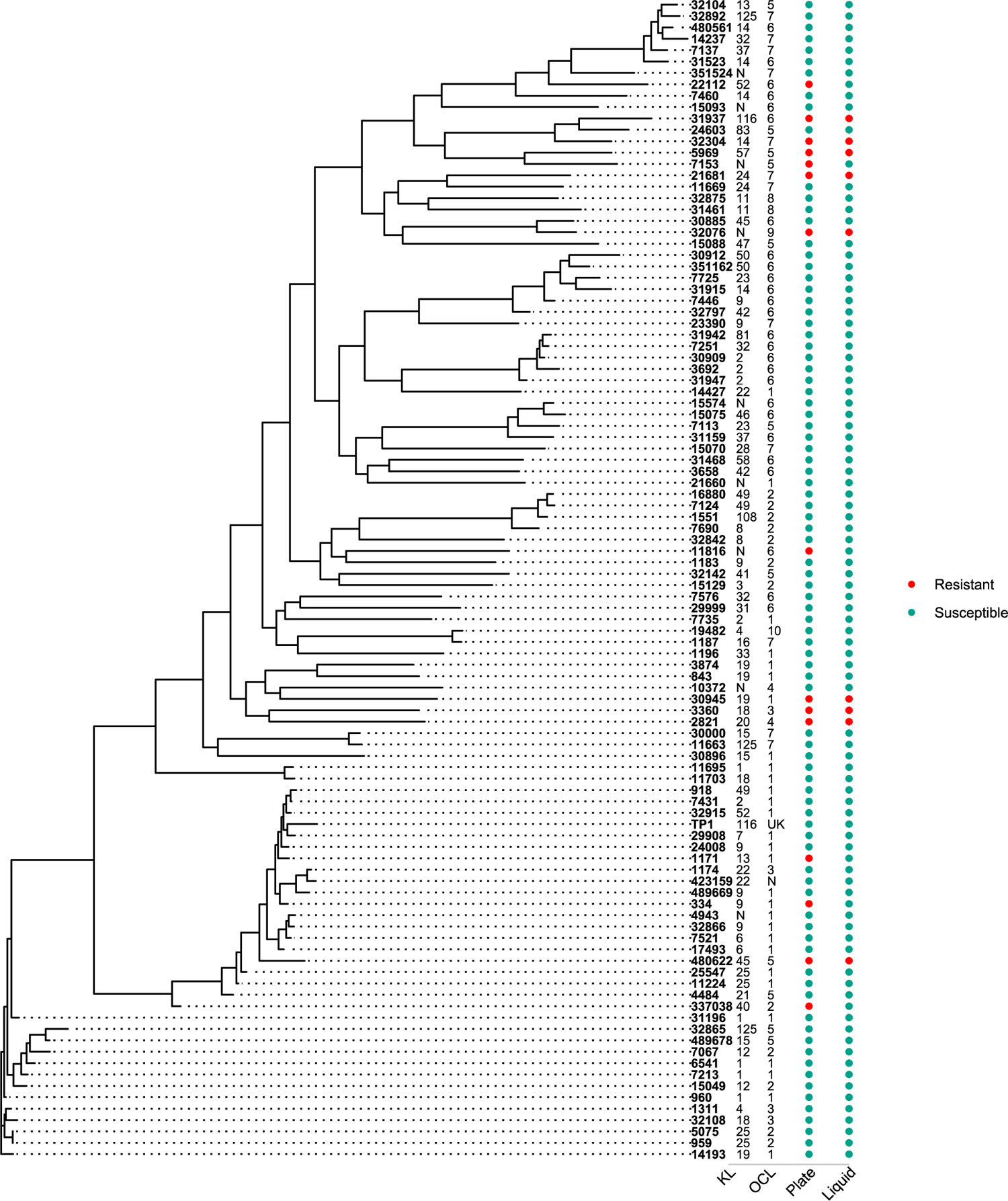
Phylogenetic tree of strains that Mystique was tested on.

Phylogenetic tree of *A. baumannii* stains that are susceptible or resistant to phage Mystique after having been tested using both plaque (labelled as ‘plate’) and liquid assays (labelled as ‘liquid’). KL indicates the strain’s capsule serotype, while OCL is the lipooligosaccharide outer core locus (N = novel and UK = unknown).

### Mystique likely targets the *A. baumannii* capsule for infection

Following the host range assay, we set out to determine the phage Mystique receptor. Due to its broad host range, we anticipated that the receptor must be something all strains have in common. This, combined with the knowledge that most other *A. baumannii* phages use the capsule as a receptor [29–31], we hypothesised that the bacterial capsule itself functions as the Mystique phage receptor. To test this, we used mutants of various capsule synthesis genes, specifically the *itrA*, *wza*, *wzb*, and *wzc* genes that encode important components of the capsular polysaccharide synthesis pathway [48] (Fig 6A). For example, ItrA is the initiating transferase, which is required for both capsule synthesis and protein O-glycosylation, while Wza, Wzb, and Wzc together form a complex that coordinates the assembly and export of the capsular polysaccharide [49,48,29].

**Figure 6.**
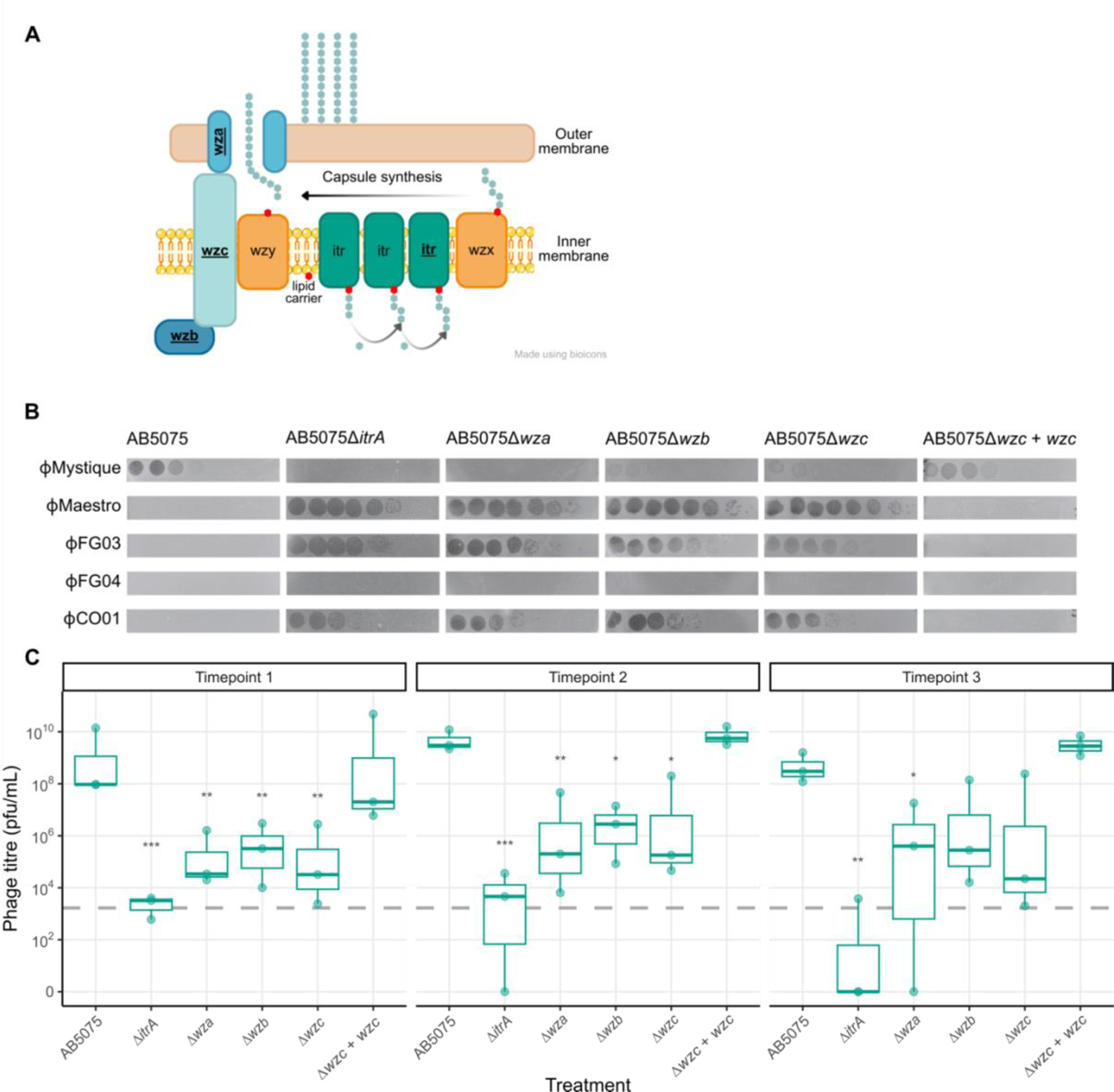
The capsule plays an important role in AB5075 susceptibility to Mystique, and susceptibility is dependent on testing environment. **A** The *A. baumannii* capsule synthesis pathway, adapted from Singh et al. 2019 [48]. **B** Serial dilution plaque assays of phages Mystique, Maestro, FG03, FG04, and CO01 on the AB5075 wild-type, Δ*itrA*::T26, Δ*wza*::T26, Δ*wzb*::T26, Δ*wzc*, and Δwzc + *wzc*. **C** Mystique phage titres over time in liquid broth culture after inoculation with the AB5075 WT or its capsule mutants. Horizontal dotted line indicates phage concentration at the start of the experiment. (Effect of treatment on phage titre; linear model, F_6,11_ = 8.44, adjusted R^2^ = 0.72: * p < 0.05, ** p < 0.01, *** p < 0.001).

Plaque assays revealed that for all four mutants there was a drastic decrease in phage Mystique infectivity (Fig 6B). Specifically, disruption of the *itrA* and *wza* genes seemed to confer complete phage resistance while we still observed some clearance on the *wzb* and *wzc* mutants, implying partial resistance (Fig 6B). We also performed plaque assay on a complemented strain: AB5075Δ*wzc* + *wzc*. Making the *wzc* gene functional again fully reversed the previously observed phage resistance, resulting in it becoming re-susceptible at the same level as the wild-type AB5075 (Fig. 6B). This further supported our hypothesis that Mystique uses the capsule itself as its receptor. Additionally, we assessed the other phages used during phage isolation on the same capsule mutants and found an inverse pattern where the phages that can amplify in liquid but not on a bacterial lawn of AB5075 (Fig 2) do cause lysis on all AB5075 capsule mutants (Fig 6B).

Finally, we tested how well Mystique would amplify on these mutants in liquid culture, following our observations on the importance of environmental conditions when assessing phage susceptibility/resistance (Figs 2B and 4). Doing this (with daily 1:100 dilutions into fresh media) revealed that Mystique can amplify in the population when inoculated with all capsule mutants, but that this was to some degree mutant dependent (Fig 6C). In particular, we observed two cases of phage extinction in the presence of the Δ*itrA* mutant specifically as well as drastically limited phage amplification, indicating this gene as being especially important in host resistance to phage Mystique. Additionally, while all mutants negatively affected Mystique amplification on day 1 and 2, by timepoint 3 there was only a significant effect of treatment (wild-type or isogenic mutant) on phage titre for the Δ*itrA* and Δ*wza* mutants, with consistently lower levels of phage for the duration of the experiment. However, across the board we never observed a complete inability of the phage to amplify as would have been expected based on the plaque assays alone. This once more made it abundantly clear how drastically environmental structure can influence observed results, and the potential for false negative interpretation of experimental findings if only conducting assays on bacterial lawns (Fig 6).

## Discussion

Here, we report on the isolation and characterisation of Mystique, a novel lytic *A. baumannii* phage – the first readily available phage against the wild-type of the clinical model strain AB5075 – as well as underscoring the limitations of conventional phage isolation techniques. Mystique is a double stranded DNA phage with a T=9 icosahedral head and helical C6 symmetric tail structure, and is the first phage to be isolated to specifically target the clinical model strain AB5075. While genetically similar to other *A. baumannii* phages, cryoEM revealed that Mystique was more structurally similar to other phages, such as YSD1 that infects *Salmonella* [46]. We speculate that the particularly long phage tail may also be beneficial when binding to a thicker capsule, such as the capsulated AB5075, but more work on the detailed structure of the tail tip and receptor pair is needed for this to become clear.

In addition to targeting AB5075, Mystique has a remarkably broad host range, being able to infect 88 (85.4%) of 103 highly diverse *A. baumannii* strains based on plaque assays. During the phage isolation process, however, we also consistently observed how some phages are able to infect AB5075 in liquid culture while simultaneously not plaquing on bacterial lawns (Fig 2), the latter of which is the standard method for isolating phages and testing infectivity [36]. When also assessing Mystique in liquid culture, its final host range was at 94 (91.3%) out of the 103 strains tested.

The broad host range observed is likely due to Mystique using the *A. baumannii* capsule as its receptor, which is a common receptor for other *A. baumannii* phages as well. Interestingly, most phages seem restricted by capsule serotype [29] in a way Mystique is not. The capsule being the phage receptor was supported by how several AB5075 capsule mutants conferred either complete resistance or drastically reduced infectivity compared to the wild-type using plaque assays (Fig 6B). Yet we still found that Mystique was able to amplify, although to a lesser extent, on all capsule mutants when tested in liquid (Fig 6C). Making the *itrA* gene non-functional resulted in the strongest effect, with two out of three populations showing phage extinction by timepoint 3. Nonetheless, this ability of the phage to persist and amplify on seemingly resistant bacteria may increase the likelihood of escape phages (phages with mutational changes that overcome host resistance) to emerge over time[50].

Overall, this disconnect between the standard assays on bacterial lawn and liquid culture likely means we are missing a fundamental property when it comes to phages against *A. baumannii* and potentially also other bacteria with similar ability to regulate capsule thickness. For instance, *E. coli* was recently found to regulate capsule thickness and consequent masking of the phage receptor in response to cell surface pressure with downstream effects on phage susceptibility [51]. This effect was lost if using an *E. coli* Δ*wza* mutant [51], which we note is one of the same genes disrupted in our work. It is therefore not unlikely that a similar effect may be involved for *A. baumannii*, and more work should be done to elucidate what causes some phages to be unable to infect on bacterial lawns but not in liquid environments.

This imperfect mapping between testing environments further highlights the complex nature of bacteria-phage dynamics and the need for research on the finer mechanistic details at play when phages use the *A. baumannii* capsule as their receptor. This was made clear by our results showing how multiple phages in our collection that use the capsule as their receptor[20,30] were also only able to lyse AB5075 capsule negative mutants. Overall, this indicates that we are still missing crucial pieces of the puzzle regarding how various phages interact with a diverse and plastic surface structure like the *A. baumannii* capsule [26,27,48].

In conclusion, the Mystique phage isolation, structural analysis, and characterisation process highlights the importance of re-evaluating traditional phage isolation techniques and adopting a multifaceted approach to phage research. By interrogating the interplay between phages and their bacterial hosts in diverse environmental contexts we can gain deeper insights into the mechanisms of phage resistance in ways that will aid us in devising more robust strategies for phage therapy against *A. baumannii* and other bacteria with complex capsules.

## Materials and Methods

### Bacterial strains and phages

The strain used for isolating phage was *A. baumannii* AB5075_UW [25]. An additional 102 *A. baumannii* strains were used to assess phage host range. These included 100 diverse clinical isolates from the Multidrug-Resistant Organism Repository and Surveillance Network (MRSN) [34], clinical isolate FZ21 from Queen Astrid Military Hospital, Belgium [17], and clinical isolate TP1 from UC San Diego, USA [20]. The capsule mutants *wza*::T26, *Δwzb*:T26, and Δ*itrA*::T26) were obtained from the AB5075 transposon mutant library [52] while the *Δwzc* mutant and complemented Δ*wzc* + *wzc* are both previously described [53]. The additional phages used for this study were Maestro [20], FG03, FG04, and CO01 [30].

### Phage isolation

Mystique was isolated from raw sewage water from the R.L. Sutton Water Reclamation Facility in Atlanta, USA. Methods for isolation were adapted from previously used methods [31]. In brief, for a final concentration of 3 g of powdered LB medium (VWR) was mixed with 100 mL of raw sewage water before 100 µL of AB5075 was added. Bacteria were grown to exponential phase before being added the sewage, after which they were incubated in the sewage mixture overnight at 37°C at 180rpm.

After inoculation overnight, 1 mL of the sewage/LB mixture was sampled and centrifuged for 5 minutes at 8000 x g before the supernatant was filtered through a 0.22µM spin-X centrifuge tube filter (Corning) at 6000 x g to remove any remaining bacterial cells. 10 µL of this filtrate was added to 100 µL of AB5075 in exponential growth phase before incubation for 20 minutes at 37°C and 180 rpm. After this second round of inoculation, the 100 µL mixture was combined with 2.5 mL of top agar (0.5% LB agar, VWR) before being poured over LB agar plates and placed in an incubator at 37°C overnight. This, however, yielded no phage plaques and so 100 µL of supernatant from the raw sewage water/LB powder mixture was added to 6 mL of LB medium with AB5075 at exponential growth. In addition to the sewage filtrate, other known *A. baumannii* phages were added to the mixture in an attempt at limiting the rapid evolution of phage resistance overnight. These phages were Maestro [20], FG03, FG04, and CO01 [30]. These cultures were subsequently grown overnight, before taking 1mL of the culture to be centrifuged at 8000 x g for 5 minutes and filtering the resulting supernatant through a 0.22 µM filter. 5 µL of the filtrate was then pipetted on top of a lawn of AB5075 before incubation at 37°C overnight. This resulted in bacterial clearance, and a 1-10 µL pipette tip was used to transfer a small amount from the centre of the zone of clearance into a fresh bacterial culture of AB5075. This was done three times, but no individual phage plaques were seen on AB5075.

To ensure the isolate only contained one phage, the lysate after three days of passaging and purification was also tested on an *A. baumannii* host for some of the other phages initially added in the cocktail: MRSN strain 423159 [34]. Clear individual phage plaques were observed on 423159 from which one was picked and purified three times (repeated plaque assays on 423159 followed by inoculation with AB5075).

### Phage sequencing, annotation, and assembly

Phage DNA was extracted following already established methods for extracting phage DNA[54]. In short: 500µL of filter-sterilised phage lysate was incubated statically with 50 µL DNase I 10x buffer, 1 µL DNase I (1 U/µL), and 1 µL RNase A (10 mg/mL) for 1.5 h at 37 °C. Following this step, 0.5 M EDTA was added for a final concentration of 20 mM before 1.25 µL of Proteinase K was added after which the sample was inoculated at 56°C for another 1.5 h. After this second incubation step, DNA was extracted using following the instruction of the DNeasy Blood and Tissue Kit (Qiagen).

Following extraction, DNA fragmentation was performed using the NEBNext® Ultra™ II FS DNA Library Prep Kit (New England Biolabs), an enzymatic fragmentation assay with an average fragment size of 380 bp. After fragmentation, the fragmented DNA was end-repaired, A-tailed, and ligated with Illumina-compatible adaptors using the same NEBNext® kit. The ligated products were then amplified via PCR to enrich the library. The amplified libraries were purified using AMPure XP beads (Beckman Coulter) to remove any unbound adaptors and smaller fragments.

The prepared libraries were evaluated for size distribution and concentration using the Agilent 2100 Bioanalyzer (Agilent Technologies) with a High Sensitivity DNA Kit. Libraries exhibiting the desired size range and absence of primer-primer dimers were selected for sequencing on the Illumina NovaSeq 5000 platform, employing a paired-end 150 bp (PE150) read configuration to generate high-quality short reads.

For long-read sequencing, the extracted DNA was prepared for sequencing using the Ligation Sequencing Kit (SQK-LSK109) from Oxford Nanopore Technologies (ONT, Oxford, UK). The extracted DNA was quantified using a Qubit 4 Fluorometer (Thermo Fisher Scientific) to ensure an adequate amount of input material. The DNA was subjected to end-repair and dA-tailing using the NEBNext® Ultra™ II End Repair/dA-Tailing Module (New England Biolabs). Following end-repair and dA-tailing, ONT’s proprietary sequencing adaptors were ligated to the DNA fragments using the Blunt/TA Ligase Master Mix (New England Biolabs) provided in the Ligation Sequencing Kit. The ligation reaction mixture was purified using AMPure XP beads (Beckman Coulter) to remove unligated adaptors and small DNA fragments, ensuring that only high-molecular-weight DNA with ligated adaptors proceeded to sequencing. The purified library was quantified again using the Qubit 4 Fluorometer to confirm the concentration and ensure that an adequate amount of library was available for sequencing. The prepared library was loaded onto a Flow Cell (R9.4.1) and sequenced on the Oxford Nanopore MinION device. Sequencing was performed according to the manufacturer’s standard operating procedures, and run conditions were monitored using ONT’s MinKNOW software. Sequencing continued until sufficient data was generated to achieve the desired genome coverage.

The sequencing data from both platforms were processed and analysed using standard bioinformatics pipelines. Short reads from the Illumina platform were trimmed and assembled using SPAdes, while long reads from the Nanopore platform were basecalled using Guppy and assembled using Canu.

Following sequencing, hybrid genome assembly and annotation were conducted on the Galaxy [55] and Web Apollo [56] phage annotation platforms. Unless otherwise noted, default parameters were used for all software. Long reads under 1kb were filtered out using Filtlong v.0.1.2 [57] and were subsequently quality checked using Nanoplot v.1.41.0 [58,59]. Flye v.2.9.1 [60,61] was used with –nano-hq and metagenomic assembly parameters [62] to obtain a consensus draft assembly. One circular contig 77,172bp in length with 3,257x coverage was obtained. Short sequencing reads were rarefied to 100x coverage to improve assembly quality using FastQ Subset v.1.1 [63,64] and trimmed using the Trim Sequences tool v.1.0.2 [65]. Short reads were quality checked using FastQC v.0.72 [66] and aligned with the long read draft assembly using the Map with BWA-MEM tool v.0.7.17.2 [67,68]. That output was then used with pilon v.1.20.1 [69] to create a consensus hybrid assembly. The complete assembled contig was 77,401 bp long and was reopened using Reopen Fasta Sequences v.2.0 [64] in order to avoid interrupting genes. BLASTn [70] was used to find similarity to previously-identified phages.

The final Mystique assembly was imported into Apollo using the Galaxy Structural Phage Annotation Workflow v.2023.01 and the locations of genes were predicted as described in Ramsey *et al*. [63] using GLIMMER3 v.0.2 [71], MetaGeneAnnotator v.1.0.0 [72], and SixPack v.5.0.0 [73]. The criteria weighed in order to manually make final gene calls were assessment of gaps and overlaps between genes, the presence of a valid Shine-Dalgarno sequence, and the presence of a valid start codon. The presence of tRNAs was assessed using tRNAScan-SE v.0.4 [74] and ARAGORN v.0.6 [75]. When structural annotation was complete, functional annotation was conducted using the Galaxy Functional Phage Annotation Workflow v.2023.01 [63]. BLASTp [76,77] results were compared from the canonical phages, nonredundant-all phages, and Swiss-Prot databases to manually annotate putative functions. The geecee tool v.5.0.0 [78] was used to determine the GC content of the Mystique genome.

### Transmission electron microscopy and cryogenic electron microscopy

The Mystique phage sample was prepared using PEG precipitation[40] before being added to a plasma cleaned continuous carbon grid (30s, hydrogen, oxygen) and stained with 2% uranyl formate. The grids were imaged on a Hitachi 7800 TEM operated at 100kV, and data were collected at a pixel size of 1.77Å with a TVIPS XF416 (Gatan).

The phage sample was further concentrated 50-fold using Amicon Ultra 100k concentrators (100k MWCO). Grids were frozen on Quantifoil R2/2 Cu 300 mesh grids using a Vitrobot Mark IV (ThermoFisher Scientific) at 20°C and 100% humidity with a wait time of 0s, a blot time of 6.5s and a blot force of 1. The grids were then clipped into Autogrids and imaged on a Titan Krios G2 (ThermoFisher Scientific) equipped with a Gatan K3 direct electron detector and BioQuantum energy filter set to 20 eV slit width. Data were collected at a pixel size of 1.083Å, a dose of 49.90 e^-^/Å^2^, and a nominal defocus range of −1 to −2 µm. With fringe-free imaging (FFI), we were able to collect 6 images per hole, totalling in 8565 images for the data collection. Data were acquired using Leginon [79,80] on NYSBC Krios1, dataset m23oct30a.

All data were processed in cryoSPARC v4 [81] using standard workflow starting with raw frames. Frames were imported and motion-corrected using patch motion job and CTF was estimated using patch CTF job. Micrographs were sorted to exclude any with CTF estimation >4.6 Å. For both heads and tails, particles were first manually picked, then initial 2D classification was used to generate templates. For heads template particle picking was used resulting in ∼26,000 particles picked that were triaged by 2D classification to yield a subset of 6000 particles. Initial models were generated using *ab-initio* jobs and resulting models were used for homogeneous refinement with icosahedral symmetry applied. For tails we used filament tracer and initially extracted ∼1 million segments that were triaged by 2D classification to 191,000 particles. To generate an initial model first ab-initio job was used, followed by homogeneous refinement. The resulting map was used to identify initial helical parameters and was also used as an initial model for helical refinement. After initial helical refinement in a second round C6 symmetry was enforced.

For *de novo* structure prediction of the tail we used ModelAngelo with a no_seq flag [44]. The resulting prediction was used to first identify the gene in the Mystique genome using NCBI tBLASTn. The identified protein was manually built into the tail density using COOT[82] and further refined with *Phenix* [83]. A structural similarity search was done with Foldseek with default parameters. The structure prediction of Mystique head major capsid protein was done using AlphaFold2 [84] collab notebook using the sequence annotated in Mystique genome. CryoEM maps of an empty and full head are uploaded to EMDB with the following codes: EMD (pending). Tail model was deposited to PDB (pending).

#### *A. baumannii* phylogenetic analysis

Host genomes were downloaded from NCBI, accessed in May of 2024. A maximum parsimony tree based on whole-genome single nucleotide polymorphisms (SNPs) was constructed using kSNP4.1 [85], with a Kmer size of 17. The tree was rooted on strain 11669 (accession number GCF_006493685.1), as described in Galac *et al*. 2022 [34]. The resulting tree was visualized in R using ggtree v3.12.0 [86] and ggtreeExtra v1.14.0 [87]. To generate a chronogram from the phylogram, we utilized the chronos function in ape v5.0 [88]. We tested all available models (correlated, discrete, relaxed, and clock), and found that the tree generated under a strict clock was favoured. Using the chronogram, we calculated the phylogenetic D statistic [47] using function phylo.d from caper v1.0.3 [89].

### Testing of phage host range and receptor

The initial experiment to test Mystique host range were done using plaque assays. These plaque assays on 103 highly diverse clinical *A. baumannii* strains were done by growing the bacterial strains overnight in glass microcosms containing 6 mL of LB broth while shaking at 180 rpm at 37°C. The following day, 200 µl of each individual bacterial culture was mixed with 10 mL of 0.5% LB agar before gentle mixing and pouring on top of LB agar plates. This top layer was left to dry for approximately 30 minutes, followed by pipetting 5 µL of serial diluted phage Mystique on top of the dried top agar layer (*n* = 3 per plate). 1:10 serial dilutions of Mystique were prepared in 96-well plates, with approximately 1 x 10^9^ pfu/mL as the undiluted phage concentration. These plates were then incubated overnight at 37°C before being checked for phage clearance the following day.

All phage host range and infectivity experiments in liquid were done by inoculating 60 µL from overnight cultures of *A. baumannii* strains into glass microcosms containing 6 mL of LB medium. 10^4^ pfu/mL of phage Mystique (Figs 1, 6, and S3) or Maestro, FG03, FG04, or CO01 (Fig 2) were then added to the glass microcosms followed by incubation overnight at 37°C at 180 rpm (*n* = 3 per treatment). Transfers of 1:100 into fresh LB were done daily for a total of three days, and phage titres were either tracked daily (Fig 6) or assessed by the end of the experiment (Figs 1, 2, and S3) by pipetting serial dilutions of chlorophorm-treated lysate on lawns of *A. baumannii* 43159.

Experiments to test for the phage receptor was done using plaque assays of phage Mystique on lawns of *Δwza*::T26, *Δwzb*:T26, *Δwzc*, and Δ*itrA*::T26 as well as Δ*wzc* + *wzc* and the AB5075 wild-type. Additionally, an evolution experiment tracking phage titres over time when inoculated with all isogenic mutants and the wild-type strain was done as described above. All data is available at (pending): 10.6084/m9.figshare.26180200

## Acknowledgements

The authors would like to thank David Pride for providing us with *A. baumannii* strain TP1, Ry Young for phage Maestro, and Jeremy Barr for phages FG03, FG04, and CO01. EOA would like to thank her mentors Marvin Whiteley, Steve Diggle, Sam Brown, and Brian Hammer for support and feedback on experimental design and the final manuscript. Funding for this research was provided to EOA by the Center for Microbial Dynamics and Infection’s Early Career Award Fellow award. The structural analyses by MK and DB were performed at the Simons Electron Microscopy Center at the New York Structural Biology Center, with major support from the Simons Foundation (SF349247). PNR is supported by Department of Veterans Affairs grants I01BX001725 and IK6BX004470 (Senior Research Career Scientist award).

---

**Supplemental figure 1. Structure prediction of Mystique’s major capsid protein structure suggests HK97 fold. A** AlphaFold2 structure prediction shows major structural features of HK97 fold: A- and P-domains with a characteristic backbone helix and E-loop. **B** Rigid-body fitting of the predicted structure into the experimental density forming an asymmetric unit. **C** Icosahedral symmetrisation of the asymmetric unit fills most of the capsid’s density, while unfilled densities located at trifold or pseudotrifold locations suggest an unidentified cement or decoration protein.

**Supplemental figure 2. Structure of a Mystique’s tail monomer is highly similar to a phlagellotropic bacteriophage YSD1. A** Side by side comparison of a YSD1 tail protein monomer (top, yellow) and Mystique phage tail protein monomer (bottom, blue). These proteins appear structurally very similar yet share very low primary sequence similarity as shown by the **B** sequence alignment of YSD1 tail protein (upper sequence) and Mystique tail protein (lower sequence). Additionally, Mystique’s tail protein lacks a C-terminal domain and has a truncated N-terminal domain. **C** Cross-section view of the tail model where residues were coloured by their electronegativity showing the negatively charged central cavity.

Supplemental figure 3. Mystique can infect and amplify on strains in liquid that it does not lyse on a lawn. Out of the 15 strains Mystique was unable to lyse on a bacterial lawn, six proved to be susceptible in liquid culture.

**Supplemental figure 4. Testing for phylogenetic signal across Mystique susceptible strains.** D values (black vertical lines) as a measure of phylogenetic signal, where a D value of 1 (red lines) indicates randomness and 0 (blue lines) implies departure from the randomness expected under a Brownian evolution threshold model. Calculated for *A. baumannii* strains susceptible to phage Mystique either on **A** plate or in **B** liquid (Fig. 5).

## References

1. Antunes LCS, Visca P, Towner KJ. *Acinetobacter baumannii*: evolution of a global pathogen. Pathog Dis. 2014;71: 292–301. doi:10.1111/2049-632X.12125

2. Cain AK, Hamidian M. Portrait of a killer: Uncovering resistance mechanisms and global spread of *Acinetobacter baumannii*. PLOS Pathog. 2023;19: e1011520. doi:10.1371/journal.ppat.1011520

3. Perez F, Hujer AM, Hujer KM, Decker BK, Rather PN, Bonomo RA. Global Challenge of Multidrug-Resistant *Acinetobacter baumannii*. Antimicrob Agents Chemother. 2007;51: 3471–3484. doi:10.1128/aac.01464-06

4. Zin EB-BR, Towner KJ. *Acinetobacter* spp. as Nosocomial Pathogens: Microbiological, Clinical, and Epidemiological Features. CLIN MICROBIOL REV. 1996;9. doi:10.1128/cmr.9.2.148

5. Centers for Disease Control and Prevention. Antibiotic Resistance Threats in the United States, 2013. Atlanta, Georgia: U.S. Department of Health and Human Services, CDC; 2013. Available: https://www.cdc.gov/drugresistance/pdf/ar-threats-2013-508.pdf

6. Ibrahim S, Al-Saryi N, Al-Kadmy IMS, Aziz SN. Multidrug-resistant *Acinetobacter baumannii* as an emerging concern in hospitals. Mol Biol Rep. 2021;48: 6987– 6998. doi:10.1007/s11033-021-06690-6

7. Lei J, Han S, Wu W, Wang X, Xu J, Han L. Extensively drug-resistant *Acinetobacter baumannii* outbreak cross-transmitted in an intensive care unit and respiratory intensive care unit. Am J Infect Control. 2016;44: 1280–1284. doi:10.1016/j.ajic.2016.03.041

8. Aygün G, Demirkiran O, Utku T, Mete B, Ürkmez S, Yılmaz M, et al. Environmental contamination during a carbapenem-resistant *Acinetobacter baumannii* outbreak in an intensive care unit. J Hosp Infect. 2002;52: 259–262. doi:10.1053/jhin.2002.1300

9. Bernards AT, Harinck HIJ, Dijkshoorn L, van der Reijden TJK, van den Broek PJ. Persistent *Acinetobacter baumannii*? Look Inside Your Medical Equipment. Infect Control Hosp Epidemiol. 2004;25: 1002–1004. doi:10.1086/502335

10. Cruz-López F, Martínez-Meléndez A, Villarreal-Treviño L, Morfín-Otero R, Maldonado-Garza H, Garza-González E. Contamination of healthcare environment by carbapenem-resistant *Acinetobacter baumannii*. Am J Med Sci. 2022;364: 685–694. doi:10.1016/j.amjms.2022.07.003

11. Wilks M, Wilson A, Warwick S, Price E, Kennedy D, Ely A, et al. Control of an Outbreak of Multidrug-Resistant *Acinetobacter baumannii-calcoaceticus* Colonization and Infection in an Intensive Care Unit (ICU) Without Closing the ICU or Placing Patients in Isolation. Infect Control Hosp Epidemiol. 2006;27: 654– 658. doi:10.1086/507011

12. LaVergne S, Hamilton T, Biswas B, Kumaraswamy M, Schooley RT, Wooten D. Phage Therapy for a Multidrug-Resistant *Acinetobacter baumannii* Craniectomy Site Infection. Open Forum Infect Dis. 2018;5: ofy064. doi:10.1093/ofid/ofy064

13. Schooley RT, Biswas B, Gill JJ, Hernandez-Morales A, Lancaster J, Lessor L, et al. Development and Use of Personalized Bacteriophage-Based Therapeutic Cocktails To Treat a Patient with a Disseminated Resistant *Acinetobacter baumannii* Infection. Antimicrob Agents Chemother. 2017;61: e00954–17, e00954-17. doi:10.1128/AAC.00954-17

14. Jessup CM, Kassen R, Forde SE, Kerr B, Buckling A, Rainey PB, et al. Big questions, small worlds: microbial model systems in ecology. Trends Ecol Evol. 2004;19: 189–197. doi:10.1016/j.tree.2004.01.008

15. McDonald MJ. Microbial Experimental Evolution – a proving ground for evolutionary theory and a tool for discovery. EMBO Rep. 2019;20: e46992. doi:10.15252/embr.201846992

16. Conners R, León-Quezada RI, McLaren M, Bennett NJ, Daum B, Rakonjac J, et al. Cryo-electron microscopy of the f1 filamentous phage reveals insights into viral infection and assembly. Nat Commun. 2023;14: 2724. doi:10.1038/s41467-023-37915-w

17. Alseth EO, Pursey E, Luján AM, McLeod I, Rollie C, Westra ER. Bacterial biodiversity drives the evolution of CRISPR-based phage resistance. Nature. 2019;574: 549–552. doi:10.1038/s41586-019-1662-9

18. Guillemet M, Chabas H, Nicot A, Gatchich F, Ortega-Abboud E, Buus C, et al. Competition and coevolution drive the evolution and the diversification of CRISPR immunity. Nat Ecol Evol. 2022; 1–9. doi:10.1038/s41559-022-01841-9

19. Romeyer Dherbey J, Bertels F. The untapped potential of phage model systems as therapeutic agents. Virus Evol. 2024;10: veae007. doi:10.1093/ve/veae007

20. Liu M, Hernandez-Morales A, Clark J, Le T, Biswas B, Bishop-Lilly KA, et al. Comparative genomics of *Acinetobacter baumannii* and therapeutic bacteriophages from a patient undergoing phage therapy. Nat Commun. 2022;13: 3776. doi:10.1038/s41467-022-31455-5

21. Bull JJ, Molineux IJ. Predicting evolution from genomics: experimental evolution of bacteriophage T7. Heredity. 2008;100: 453–463. doi:10.1038/sj.hdy.6801087

22. Wichman HA, Brown CJ. Experimental evolution of viruses: Microviridae as a model system. Philos Trans R Soc B Biol Sci. 2010;365: 2495–2501. doi:10.1098/rstb.2010.0053

23. Budzik JM, Rosche WA, Rietsch A, O’Toole GA. Isolation and Characterization of a Generalized Transducing Phage for *Pseudomonas aeruginosa* Strains PAO1 and PA14. J Bacteriol. 2004;186: 3270–3273. doi:10.1128/JB.186.10.3270-3273.2004

24. Cady KC, Bondy-Denomy J, Heussler GE, Davidson AR, O’Toole GA. The CRISPR/Cas Adaptive Immune System of *Pseudomonas aeruginosa* Mediates Resistance to Naturally Occurring and Engineered Phages. J Bacteriol. 2012;194: 5728–5738. doi:10.1128/JB.01184-12

25. Jacobs AC, Thompson MG, Black CC, Kessler JL, Clark LP, McQueary CN, et al. AB5075, a Highly Virulent Isolate of Acinetobacter baumannii, as a Model Strain for the Evaluation of Pathogenesis and Antimicrobial Treatments. mBio. 2014;5: 10.1128/mbio.01076-14.

26. Tipton KA, Dimitrova D, Rather PN. Phase-Variable Control of Multiple Phenotypes in *Acinetobacter baumannii* Strain AB5075. J Bacteriol. 2015;197: 2593–2599. doi:10.1128/JB.00188-15

27. Chin CY, Tipton KA, Farokhyfar M, Burd EM, Weiss DS, Rather PN. A high-frequency phenotypic switch links bacterial virulence and environmental survival in *Acinetobacter baumannii*. Nat Microbiol. 2018;3: 563–569. doi:10.1038/s41564-018-0151-5

28. Pérez-Varela M, Singh R, Colquhoun JM, Starich OG, Tierney ARP, Tipton KA, et al. Evidence for Rho-dependent control of a virulence switch in *Acinetobacter baumannii*. Biswas I, editor. mBio. 2023; e02708–23. doi:10.1128/mbio.02708-23

29. Bai J, Raustad N, Denoncourt J, van Opijnen T, Geisinger E. Genome-wide phage susceptibility analysis in *Acinetobacter baumannii* reveals capsule modulation strategies that determine phage infectivity. PLOS Pathog. 2023;19: e1010928. doi:10.1371/journal.ppat.1010928

30. Gordillo Altamirano F, Forsyth JH, Patwa R, Kostoulias X, Trim M, Subedi D, et al. Bacteriophage-resistant *Acinetobacter baumannii* are resensitized to antimicrobials. Nat Microbiol. 2021; 1–5. doi:10.1038/s41564-020-00830-7

31. Regeimbal JM, Jacobs AC, Corey BW, Henry MS, Thompson MG, Pavlicek RL, et al. Personalized Therapeutic Cocktail of Wild Environmental Phages Rescues Mice from *Acinetobacter baumannii* Wound Infections. Antimicrob Agents Chemother. 2016;60: 5806–5816. doi:10.1128/AAC.02877-15

32. Peters DL, Davis CM, Harris G, Zhou H, Rather PN, Hrapovic S, et al. Characterization of Virulent T4-Like *Acinetobacter baumannii* Bacteriophages DLP1 and DLP2. Viruses. 2023;15: 739. doi:10.3390/v15030739

33. Scholl D, Adhya S, Merril C. *Escherichia coli* K1’s Capsule Is a Barrier to Bacteriophage T7. Appl Environ Microbiol. 2005;71: 4872–4874. doi:10.1128/AEM.71.8.4872-4874.2005

34. Galac MR, Snesrud E, Lebreton F, Stam J, Julius M, Ong AC, et al. A Diverse Panel of Clinical *Acinetobacter baumannii* for Research and Development. Antimicrob Agents Chemother. 2020;64: e00840–20. doi:10.1128/AAC.00840-20

35. d’Herelle F. An invisible microbe that is antagonistic to the dysentery bacillus. Comptes Rendus Acad Sci. 1917;165: 373–375.

36. van Charante F, Holtappels D, Blasdel B, Burrowes B. Isolation of Bacteriophages. In: Harper DR, Abedon ST, Burrowes BH, McConville ML, editors. Bacteriophages: Biology, Technology, Therapy. Cham: Springer International Publishing; 2019. pp. 1–32. doi:10.1007/978-3-319-40598-8_14-1

37. Mardiana M, Teh S-H, Lin L-C, Lin N-T. Isolation and Characterization of a Novel Siphoviridae Phage, vB_AbaS_TCUP2199, Infecting Multidrug-Resistant Acinetobacter baumannii. Viruses. 2022;14: 1240. doi:10.3390/v14061240

38. Margulieux KR, Bird JT, Kevorkian RT, Ellison DW, Nikolich MP, Mzhavia N, et al. Complete genome sequence of the broad host range *Acinetobacter baumannii* phage EAb13. Microbiol Resour Announc. 2023;12: e00341–23. doi:10.1128/MRA.00341-23

39. Conant GC, Wolfe KH. GenomeVx: simple web-based creation of editable circular chromosome maps. Bioinformatics. 2008;24: 861–862. doi:10.1093/bioinformatics/btm598

40. Luong T, Salabarria A-C, Edwards RA, Roach DR. Standardized bacteriophage purification for personalized phage therapy. Nat Protoc. 2020;15: 2867–2890. doi:10.1038/s41596-020-0346-0

41. Carroll-Portillo A, Coffman CN, Varga MG, Alcock J, Singh SB, Lin HC. Standard Bacteriophage Purification Procedures Cause Loss in Numbers and Activity. Viruses. 2021;13: 328. doi:10.3390/v13020328

42. Caspar DLD, Klug A. Physical Principles in the Construction of Regular Viruses. Cold Spring Harb Symp Quant Biol. 1962;27: 1–24. doi:10.1101/SQB.1962.027.001.005

43. Suhanovsky MM, Teschke CM. Nature׳s favorite building block: Deciphering folding and capsid assembly of proteins with the HK97-fold. Virology. 2015;479– 480: 487–497. doi:10.1016/j.virol.2015.02.055

44. Jamali K, Käll L, Zhang R, Brown A, Kimanius D, Scheres SHW. Automated model building and protein identification in cryo-EM maps. Nature. 2024;628: 450–457. doi:10.1038/s41586-024-07215-4

45. van Kempen M, Kim SS, Tumescheit C, Mirdita M, Lee J, Gilchrist CLM, et al. Fast and accurate protein structure search with Foldseek. Nat Biotechnol. 2024;42: 243–246. doi:10.1038/s41587-023-01773-0

46. Hardy JM, Dunstan RA, Grinter R, Belousoff MJ, Wang J, Pickard D, et al. The architecture and stabilisation of flagellotropic tailed bacteriophages. Nat Commun. 2020;11: 3748. doi:10.1038/s41467-020-17505-w

47. Fritz SA, Purvis A. Selectivity in mammalian extinction risk and threat types: a new measure of phylogenetic signal strength in binary traits. Conserv Biol J Soc Conserv Biol. 2010;24: 1042–1051. doi:10.1111/j.1523-1739.2010.01455.x

48. Singh JK, Adams FG, Brown MH. Diversity and Function of Capsular Polysaccharide in *Acinetobacter baumannii*. Front Microbiol. 2019;9. Available: https://www.frontiersin.org/articles/10.3389/fmicb.2018.03301

49. Kenyon JJ, Hall RM. Variation in the Complex Carbohydrate Biosynthesis Loci of *Acinetobacter baumannii* Genomes. PLoS ONE. 2013;8: e62160. doi:10.1371/journal.pone.0062160

50. Hampton HG, Watson BNJ, Fineran PC. The arms race between bacteria and their phage foes. Nature. 2020;577: 327–336. doi:10.1038/s41586-019-1894-8

51. Mason G, Footer MJ, Rojas ER. Mechanosensation induces persistent bacterial growth during bacteriophage predation. mBio. 2023;14: e02766–22. doi:10.1128/mbio.02766-22

52. Gallagher LA, Ramage E, Weiss EJ, Radey M, Hayden HS, Held KG, et al. Resources for Genetic and Genomic Analysis of Emerging Pathogen *Acinetobacter baumannii*. J Bacteriol. 2015;197: 2027–2035. doi:10.1128/JB.00131-15

53. Tipton KA, Chin C-Y, Farokhyfar M, Weiss DS, Rather PN. Role of Capsule in Resistance to Disinfectants, Host Antimicrobials, and Desiccation in *Acinetobacter baumannii*. Antimicrob Agents Chemother. 2018;62: e01188–18. doi:10.1128/AAC.01188-18

54. Jakočiūnė D, Moodley A. A Rapid Bacteriophage DNA Extraction Method. Methods Protoc. 2018;1: 27. doi:10.3390/mps1030027

55. The Galaxy Community. The Galaxy platform for accessible, reproducible and collaborative biomedical analyses: 2022 update. Nucleic Acids Res. 2022;50: W345–W351. doi:10.1093/nar/gkac247

56. Lee E, Helt GA, Reese JT, Munoz-Torres MC, Childers CP, Buels RM, et al. Web Apollo: a web-based genomic annotation editing platform. Genome Biol. 2013;14: R93. doi:10.1186/gb-2013-14-8-r93

57. Wick R. rrwick/Filtlong. 2024. Available: https://github.com/rrwick/Filtlong

58. De Coster W, D’Hert S, Schultz DT, Cruts M, Van Broeckhoven C. NanoPack: visualizing and processing long-read sequencing data. Bioinformatics. 2018;34: 2666–2669. doi:10.1093/bioinformatics/bty149

59. De Coster W. wdecoster/NanoPlot. 2024. Available: https://github.com/wdecoster/NanoPlot

60. Lin Y, Yuan J, Kolmogorov M, Shen MW, Chaisson M, Pevzner PA. Assembly of long error-prone reads using de Bruijn graphs. Proc Natl Acad Sci. 2016;113: E8396–E8405. doi:10.1073/pnas.1604560113

61. Kolmogorov M. fenderglass/Flye. 2024. Available: https://github.com/fenderglass/Flye

62. Zablocki O, Michelsen M, Burris M, Solonenko N, Warwick-Dugdale J, Ghosh R, et al. VirION2: a short- and long-read sequencing and informatics workflow to study the genomic diversity of viruses in nature. PeerJ. 2021;9: e11088. doi:10.7717/peerj.11088

63. Ramsey J, Rasche H, Maughmer C, Criscione A, Mijalis E, Liu M, et al. Galaxy and Apollo as a biologist-friendly interface for high-quality cooperative phage genome annotation. PLOS Comput Biol. 2020;16: e1008214. doi:10.1371/journal.pcbi.1008214

64. TAMU-CPT/galaxy-tools. Center for Phage Technology; 2022. Available: https://github.com/TAMU-CPT/galaxy-tools

65. Gordon A. FASTQ/A short-reads pre-processing tools. 2010 [cited 20 Mar 2024]. Available: http://hannonlab.cshl.edu/fastx_toolkit/

66. Andrews S. Babraham Bioinformatics - FastQC A Quality Control tool for High Throughput Sequence Data. 2010 [cited 20 Mar 2024]. Available: https://www.bioinformatics.babraham.ac.uk/projects/fastqc/

67. Li H, Durbin R. Fast and accurate short read alignment with Burrows–Wheeler transform. Bioinformatics. 2009;25: 1754–1760. doi:10.1093/bioinformatics/btp324

68. Li H. Aligning sequence reads, clone sequences and assembly contigs with BWA-MEM. arXiv; 2013. doi:10.48550/arXiv.1303.3997

69. Walker BJ, Abeel T, Shea T, Priest M, Abouelliel A, Sakthikumar S, et al. Pilon: An Integrated Tool for Comprehensive Microbial Variant Detection and Genome Assembly Improvement. PLOS ONE. 2014;9: e112963. doi:10.1371/journal.pone.0112963

70. NCBI Resource Coordinators. Database resources of the National Center for Biotechnology Information. Nucleic Acids Res. 2018;46: D8–D13. doi:10.1093/nar/gkx1095

71. Delcher AL, Harmon D, Kasif S, White O, Salzberg SL. Improved microbial gene identification with GLIMMER. Nucleic Acids Res. 1999;27: 4636–4641. doi:10.1093/nar/27.23.4636

72. Noguchi H, Taniguchi T, Itoh T. MetaGeneAnnotator: Detecting Species-Specific Patterns of Ribosomal Binding Site for Precise Gene Prediction in Anonymous Prokaryotic and Phage Genomes. DNA Res. 2008;15: 387–396. doi:10.1093/dnares/dsn027

73. Madeira F, Park YM, Lee J, Buso N, Gur T, Madhusoodanan N, et al. The EMBL-EBI search and sequence analysis tools APIs in 2019. Nucleic Acids Res. 2019;47: W636–W641. doi:10.1093/nar/gkz268

74. Lowe TM, Chan PP. tRNAscan-SE On-line: integrating search and context for analysis of transfer RNA genes. Nucleic Acids Res. 2016;44: W54–W57. doi:10.1093/nar/gkw413

75. Laslett D, Canback B. ARAGORN, a program to detect tRNA genes and tmRNA genes in nucleotide sequences. Nucleic Acids Res. 2004;32: 11–16. doi:10.1093/nar/gkh152

76. Camacho C, Coulouris G, Avagyan V, Ma N, Papadopoulos J, Bealer K, et al. BLAST+: architecture and applications. BMC Bioinformatics. 2009;10: 421. doi:10.1186/1471-2105-10-421

77. Cock PJA, Chilton JM, Grüning B, Johnson JE, Soranzo N. NCBI BLAST+ integrated into Galaxy. GigaScience. 2015;4: 39. doi:10.1186/s13742-015-0080-7

78. Blankenberg D, Taylor J, Schenck I, He J, Zhang Y, Ghent M, et al. A framework for collaborative analysis of ENCODE data: making large-scale analyses biologist-friendly. Genome Res. 2007;17: 960–964. doi:10.1101/gr.5578007

79. Suloway C, Pulokas J, Fellmann D, Cheng A, Guerra F, Quispe J, et al. Automated molecular microscopy: The new Leginon system. J Struct Biol. 2005;151: 41–60. doi:10.1016/j.jsb.2005.03.010

80. Cheng A, Negro C, Bruhn JF, Rice WJ, Dallakyan S, Eng ET, et al. Leginon: New features and applications. Protein Sci Publ Protein Soc. 2021;30: 136–150. doi:10.1002/pro.3967

81. Punjani A, Rubinstein JL, Fleet DJ, Brubaker MA. cryoSPARC: algorithms for rapid unsupervised cryo-EM structure determination. Nat Methods. 2017;14: 290–296. doi:10.1038/nmeth.4169

82. Emsley P, Lohkamp B, Scott WG, Cowtan K. Features and development of Coot. Acta Crystallogr D Biol Crystallogr. 2010;66: 486–501. doi:10.1107/S0907444910007493

83. Liebschner D, Afonine PV, Baker ML, Bunkóczi G, Chen VB, Croll TI, et al. Macromolecular structure determination using X-rays, neutrons and electrons: recent developments in Phenix. Acta Crystallogr Sect Struct Biol. 2019;75: 861– 877. doi:10.1107/S2059798319011471

84. Jumper J, Evans R, Pritzel A, Green T, Figurnov M, Ronneberger O, et al. Highly accurate protein structure prediction with AlphaFold. Nature. 2021;596: 583–589. doi:10.1038/s41586-021-03819-2

85. Gardner SN, Slezak T, Hall BG. kSNP3.0: SNP detection and phylogenetic analysis of genomes without genome alignment or reference genome. Bioinforma Oxf Engl. 2015;31: 2877–2878. doi:10.1093/bioinformatics/btv271

86. Xu S, Li L, Luo X, Chen M, Tang W, Zhan L, et al. Ggtree: A serialized data object for visualization of a phylogenetic tree and annotation data. iMeta. 2022;1: e56. doi:10.1002/imt2.56

87. Xu S, Dai Z, Guo P, Fu X, Liu S, Zhou L, et al. ggtreeExtra: Compact Visualization of Richly Annotated Phylogenetic Data. Mol Biol Evol. 2021;38: 4039–4042. doi:10.1093/molbev/msab166

88. Paradis E, Schliep K. ape 5.0: an environment for modern phylogenetics and evolutionary analyses in R. Bioinformatics. 2019;35: 526–528. doi:10.1093/bioinformatics/bty633

89. Orme D, Freckleton R, Petzoldt T, Fritz S, Isaac N, Pearse W. caper: Comparative Analyses of Phylogenetics and Evolution in R. R package version 1.0.3. 2023 [cited 17 May 2024]. Available: https://cran.r-project.org/web/packages/caper/caper.pdf

